# Bilateral Mechanoreception Discrimination based on Bihemispheric Somatosensory Response Patterns is Associated with Proprioception and Motor Function After Stroke

**DOI:** 10.64898/2026.07.22.740075

**Authors:** Disha Gupta, Andria Farrens, Luis Garcia-Fernandez, Raymond Diaz Rojas, Vicky Chan, Joel Perry, Eric T. Wolbrecht, Dave Reinkensmeyer

## Abstract

**Objectives:** Stroke commonly impairs proprioception and motor function, yet the cortical sensory processes underlying these impairments remain poorly understood. Prior electrophysiological studies have primarily focused on the average magnitude of unilateral cortical sensory responses to vibration, potentially overlooking distributed and trial-to-trial features of sensory processing that may be functionally relevant to proprioceptive processing and motor performance. We therefore aimed to characterize bilateral cortical sensory responses to determine their relationships with proprioceptive and motor function.

**Methods:** EEG was recorded from forty-six individuals with chronic stroke during a rapid, passive, vibrotactile stimulation paradigm applied to the left and right fingertips. Somatosensory evoked potentials (SEPs) and event-related desynchronization (ERD) were quantified. Finger proprioceptive performance was assessed using a passive, robotic, finger crossing identification task, while motor function was evaluated using the Box and Block Test, Fugl-Meyer Assessment, and Nine Hole Peg Test. Associations with function were assessed using (i) unilateral sensory response magnitude at the contralateral parietal cortex and (ii) somatosensory decoder performance, defined as the accuracy with which a decoder identified the location of the stimulated hand (i.e. paretic vs. non-paretic) from combined bihemispheric response patterns. The association between these responses and proprioceptive ability and motor function was assessed. These associations were further evaluated jointly across multiple motor function measures using an exploratory analysis leveraging nonlinear dimensionality reduction and clustering.

**Results:** Vibrotactile stimulation of the paretic hand elicited ipsilesional SEP and ERD that were reduced in magnitude compared to stimulation of the non-paretic hand. Both decreased SEP magnitude and reduced sensory decoder performance were associated with greater finger proprioceptive error. Unlike unilateral responses, the somatosensory decoder’s performance was also strongly associated with motor function, explaining approximately 22.5% of the variance in motor performance. Dimensionality reduction and clustering across multiple motor assessment scores showed distinct subgroups, that showed significant differences in sensory decoding.

**Conclusions:** The hemispheric distribution and discriminability of cortical sensory responses are functionally relevant markers of sensorimotor integrity after stroke. Assessing relative lateralization of somatosensory responses for each hand, rather than the magnitude of dominant contralateral responses alone, may better capture the reliability of sensory processing after stroke, as well as the distributed cortical reorganization supporting sensorimotor function. These findings support the potential value of a novel decoding-based neurophysiological measure for sensory-driven rehabilitation, biomarker development, and patient stratification.

## Introduction

Stroke often leads to debilitating motor impairments that significantly affect quality of life. It is also frequently accompanied by somatosensory deficits (Kessner, et al., 2016, Klingner, et al., 2012, Connell et al., 2008, Carey, 2017), with prevalence reported to be as high as 57% (Bastos et al., 2025). These sensory impairments may arise indirectly–due to limb disuse following damage to motor cortical regions and descending corticospinal pathways (Huang et al., 2024, Schaechter et al., 2009), leading to subsequent weakening of sensory processing (Morioka et al., 2025)– or directly, through injury to somatosensory circuits (Kessner et al., 2019, Ingemanson et al., 2019). Indirect sensory deficits may improve with increased limb use during conventional motor rehabilitation, whereas direct sensory impairments may require targeted sensory interventions (Sasaki et al., 2025, Bolognini et al., 2016, Carey et al., 2011). However, post stroke sensory deficits have typically received less attention, with rehabilitation efforts largely focused on motor recovery due to its more visible impact on daily function (Sullivan and Hedman, 2008).

It is well understood that motor function fundamentally relies on somatosensory feedback (Amaral, 2013, Scott et al., 2016, Sainburg et al., 1995), with cutaneous mechanoreception and proprioception representing the principal somatosensory modalities supporting motor control.

Cutaneous mechanoreception uses receptors such as Pacinian corpuscles that detect dynamic change, and enables real-time force modulation during object manipulation, such as in grip force scaling and slip detection (Gentilucci et al., 1997). Proprioception involves feedback from muscle spindles and Golgi tendon organs (Sherrington, 1907; Proske & Gandevia, 2012; Proske, 2015), with substantial contributions from cutaneous mechanoreceptors that encode skin stretch as the joint moves (Cordo et al., 2011, Gandevia et al., 1983, Strzalkowski et al., 2017), providing critical information about limb position, force and coordination during movement (Farrer et al., 2003, Rosenkranz and Rothwell, 2012). Although these modalities are mediated by distinct peripheral receptors, they converge onto shared ascending dorsal-column medial lemniscus pathways that relay sensory information to the somatosensory cortex. Sensory feedback also plays a key role during recovery (Hoh and Semrau, 2025) and motor relearning in the presence of post-injury impairments which relies on somatosensory-driven error correction (Shadmehr et al., 2010) and the adaptation of alternative movement strategies (Matur and Öge, 2017, Abbruzzese and Berardelli, 2017, Zandvliet et al., 2020, Chen et al., 2018). These sensory inputs are integrated with motor control in complex ways and can be compensated by vision, for example, during object manipulation or limb positioning, making the contribution of sensory deficits to functional movement difficult to isolate and quantify.

Alternatively, the physiological integrity of sensory pathways can be assessed independently of motor control by combining peripheral sensory stimulation (such as vibrotactile or electrical stimulation) with measurement of cortical evoked responses using functional neuroimaging, magnetoencephalography (MEG), or electroencephalography (EEG). Typically, peripheral stimulation evokes responses in both somatosensory hemispheres, with the dominant response occurring in the contralateral hemisphere due to the predominantly crossed ascending sensory pathways. A smaller ipsilateral response is also observed (Iwamura et al., 1994) and is thought to be suppressed by interhemispheric inhibition (Hlushchuk et al., 2006), resulting in a characteristic lateralized response for each hand. Following stroke, injury and subsequent (mal)adaptive neuroplasticity can alter these responses, particularly in the injured (ipsilesional) hemisphere. However, it can also affect the uninjured (contralesional) hemisphere through changes in interhemispheric balance (Arya et al., 2024, Buetefisch 2015). This may reduce the distinctiveness of cortical representations evoked by each hand and thereby contribute to impaired proprioception and motor function. The relationship between these post-stroke alterations in somatosensory responses and deficits in proprioceptive ability and motor function remains poorly understood.

To better understand these relationships, we used vibration to stimulate the finger and measured (a) bihemispheric somatosensory evoked potentials (SEPs) and vibration-induced event-related desynchronization (ERD) at the sensorimotor cortex (Penfield and Boldrey, 1937, Roux et al., 2018), both established markers of sensorimotor cortical engagement (Chung et al., 2013, Rustamov et al., 2025). The vibrotactile stimulation paradigm was non-invasive, non-painful, and rapid (5-6 minutes), and required neither movement nor a perceptual response. We quantified (i) the cortical response evoked by the stimulation of the affected hand at the contralateral injured (ipsilesional) and ipsilateral uninjured (contralesional) hemisphere and (ii) the ability to decode the stimulated hand (affected versus unaffected) based on the combined bihemispheric sensorimotor cortical responses. We also measured (b) finger proprioception ability with a robot-based assessment task (the Crisscross task (Ingemanson et al., 2016), which involved passive finger movements controlled by a robotic device and active proprioceptive judgments by the participant; and (c) motor function using a widely-used clinical assessment, the Box and Block Test (Mathiowetz et al., 1985), which requires individuals to move as many small blocks as they can over a divider in 60 s. Previous studies have not examined whether bihemispheric cortical sensory decoding reflects proprioceptive impairment and motor function after stroke. Furthermore, decoding analysis has the advantage that it evaluates whether cortical sensory representations remain sufficiently stable and distinct on individual trials to identify the stimulated hand, whereas traditional SEP analyses quantify average response amplitude, that may obscure trial-to-trial variability and fluctuations in response consistency.

We hypothesized that cortical sensory responses evoked by vibrotactile stimulation of the affected hand, would be reduced at the contralateral injured hemisphere and this reduction would be associated with impaired proprioceptive ability of the affected hand, consistent with the shared ascending sensory pathways of cutaneous mechanoreception and proprioception. We further hypothesized that the bihemispheric sensory decoding would provide a complementary, and potentially more sensitive, measure of the relationship between cortical sensory processing, proprioceptive function, and motor function, capturing the distributed changes in the bihemispheric sensorimotor processing.

As an alternate, exploratory paradigm to assess the relationship between sensory responses and motor function beyond a singular clinical function score, we assess the same in relation to multiple complementary motor functional assessments, including overall impairment (Fugl-Meyer Assessment-Upper extremity), gross manual dexterity (Box and Block test), and fine finger dexterity (Nine Hole Peg test). Because these assessments capture different aspects of motor performance and differ in their dependence on somatosensory feedback, we considered them jointly as a high-dimensional representation of motor function. We then used a data-driven dimensionality reduction approach, t-distributed stochastic neighbor embedding (t-SNE) (van der Maaten and Hinton, 2008), followed by a clustering approach, to determine whether distinct motor-function subgroups emerged and whether these subgroups differed in their cortical somatosensory responses and proprioceptive function.

Identifying such subgroups may support a more individualized approach to rehabilitation by highlighting patients with underlying sensory impairments who may benefit from targeted sensory interventions (Schabrun and Hillier, 2009). We hypothesized that the association between somatosensory cortical response as well as proprioceptive ability and motor function will also be reflected across subgroups derived from the combined motor assessments.

## Methods

### Data

Data were collected in 46 people with chronic stroke during baseline assessments for a clinical trial of robotic hand training (Farrens et al., 2025 (*ArXiv:2511.00259*) ClinicalTrials.gov identifier NCT04818073). The inclusion criteria for participants with chronic stroke was age 18-85 years, unilateral upper extremity weakness due to a stroke (single or multiple site ischemic or hemorrhagic) more than 6 months ago. Exclusion criteria included a Box and Block Test score of < 3 blocks at the affected hand or a difference of < 20% between the left and right hand, severe stiffness of the arm or hand, metal implants, aphasia, major psychological problems, and concurrent participation in other studies. This data was collected with informed consent at University of California, Irvine (UCI), approved by the UCI local Institutional Review Board. All data was collected in accordance with the principles embodied in the Declaration of Helsinki and in accordance with local IRB requirements.

NIH Stroke scale (Kwah and Diong, 2014) was used to measure the level of stroke severity at the impaired hand. This 15-item impairment scale includes measuring alertness, limb strength and sensation, coordination, speech, neglect, communication ability, facial and visual field integrity. An NIHSS score of 0-4 is considered minor, 5-15 as moderate, 16-20 as moderate to severe, and 21-42 severe.

### Experiment Setup

The experiment setup involved the measurement of i) EEG-based SEPs elicited by vibrotactile stimulation at the left and right fingers, ii) proprioception assessed using a robot-based passive finger flexion-extension task, iii) motor function assessed using standardized clinical measures, including the Fugl-Meyer Assessment of the Upper Extremity (FMA-UE) (Fugl-Meyer et al., 1975; Crow and Harmeling-van der Wel, 2008, Platz et al., 2005) for overall motor impairment, the Box and Block Test (BBT) (Mathiowetz et al., 1985) for gross manual dexterity, and the Nine Hole Peg Test (NHPT) (Kellor et al., 1971; Mathiowetz et al., 1985) for fine motor dexterity.

For the vibrotactile paradigm, we preferentially stimulate Pacinian corpuscles (Fast Adapting Type II fibers) that respond to dynamic stretch at the fingers involved in recognizing fine textures, and important in object perception and grip. These mechanoreceptors located in the dermal layers, have a peak sensitivity around 250 Hz, and are most densely concentrated at the finger pads (Germann et al., 2021). The SEP response at these receptors and frequency range are also known to be near maximal (Strzalkowski et al., 2017). This vibration was unlikely to be stimulated during the relatively slow robotic movement during the Crisscross proprioceptive test.

### EEG-based Measurement of Vibrotactile Somatosensory Evoked Potentials

EEG was recorded noninvasively with a 19-channel full head headset (DSI-24, Wearable Sensing, CA) at a sampling rate of 300 Hz. The montage (Figure 1) included electrode set Fp1, Fp2, Fz, F3, F4, F7, F8, Cz, C3, C4, T7/T3, T8/T4, Pz, P3, P4, P7/T5, P8/T6, O1, O2, based on the international 10-20 EEG system (Klem et al., 1999). The ground was placed at Fpz with linked-ears reference. Data was recorded with BCI2000 software platform (Schalk et al., 2004, Mellinger and Schalk 2007).

**Figure 1:**
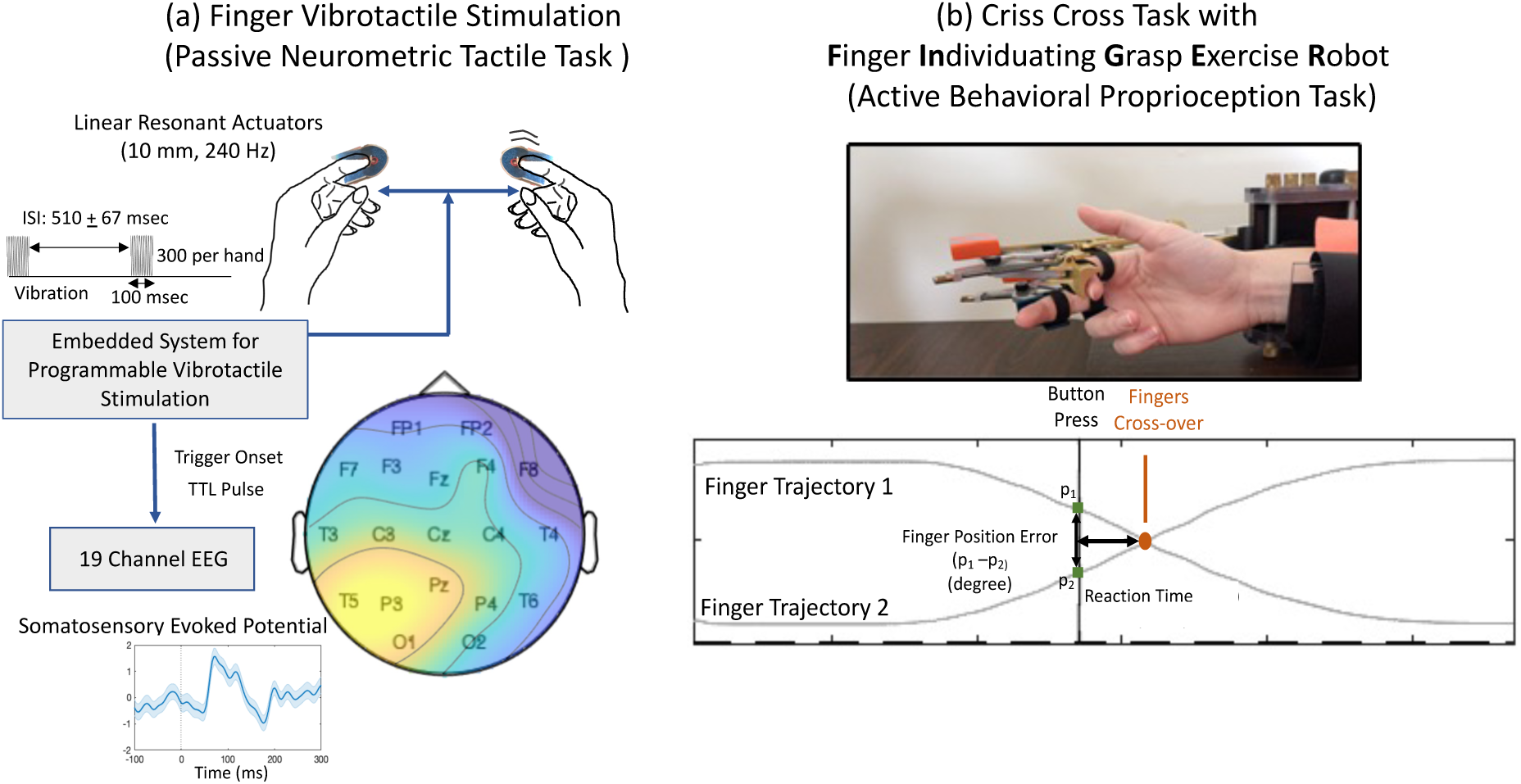
Experimental Setup: (a) Finger Vibrotactile Task: Measurement of Somatosensory Evoked Potentials (SEPs) with noninvasive electroencephalography, evoked by passive vibrotactile stimulation at left and right index finger pads. A representative SEP response to right hand stimulation is shown as a time-domain signal (evoked response) recorded over the contralateral parietal cortex. The distribution of the sensory response peak (70-90 msec) is shown through a scalp topography map. (b) Measurement of proprioception ability using the Crisscross Assessment delivered with **F**inger **In**dividuating **G**rasp **E**xercise **R**obot (FINGER): The participant is asked to indicate the moment of finger crossing while the robot moves the passive index and middle fingers in a crisscross pattern. Fingers are kept obscured from view to eliminate visual feedback. The moment of crossing is indicated by the participant pushing a button held with their other hand. The proprioception error (degree) is illustrated using an example trajectory with one cross over.

Participants were seated comfortably in an upright position with their hands resting on a soft cushion placed on the table in front of them. The cushion helped isolate the vibration and prevent it from transferring to the table and becoming audible. The vibrotactors (linear resonant actuators-8 mm diameter, resonant frequency of 240 Hz, rated operating current of 28 mA- from Precision Microdrives, UK) were positioned on the index finger pads of both hands. Small-sized splints (approximately one phalange in length) were used to maintain the position of the motor on the finger, and accommodate for spasticity and finger curling, commonly observed post-stroke. Forearms were positioned in slight supination to avoid pressure on the motors and splints. Vibrotactors were driven by an embedded system (ATmega328p microprocessor) that delivered 100 msec long vibrations per trial, with an inter stimulus interval of 400-600 msec (510 ± 67 msec). A total of 600 trials were captured, 300 per hand, randomly presented, across two consecutive runs of approximately 2.5-3 min each. Participants were instructed to fixate on a fixation cross during the run, presented on a monitor at their eye level. Prior to the assessment, vibration perception was verbally confirmed at each hand. Simultaneous to the vibration onset, the embedded system transmitted a 5V transistor-to-transistor logic (TTL) pulse to the EEG headset to mark the stimulus onsets in the EEG data. The pulses were received via an isolated digital I/O port on the EEG headset.

### Robot-based Measurement of Proprioception

Finger Individuating Grasp Exercise Robot (FINGER) (Taheri et al., 2014) was used for finger proprioception measurement with the Crisscross task (Ingemanson et al., 2019, Ingemanson et al., 2016). These measurements were made twice at baseline, in two separate sessions.

FINGER uses an 8-bar mechanism to dynamically control the orientation and position of each of the index and middle fingers individually. In the Crisscross task, FINGER passively moved the index and middle fingers continuously in a grasping motion between 12 degrees extension and 54 degrees flexion at the MCP joint, causing the fingers to cross each other 20 times. The crossings occurred at pseudorandomized speeds between 8-18 deg/sec, preventing predictability of the crossover time. The task involved sensing the moment of alignment (i.e. “crossover”) of the two fingers and pressing a button with the other hand to indicate that instance of proprioceptive perception. Finger movement was visually obscured by a screen. Finger proprioception ability was measured as the mean absolute positional error (in degrees) at button. In healthy older adults a proprioception error of 2.6 ± 6.3 degrees has been reported (Ingemanson et al., 2016).

### Motor Function Assessments

Data from three standardized tests of motor assessment was used: (i) *Fugl**-**Meyer Assessment of the Upper Extremity (FMA-UE)* (Fugl-Meyer et al., 1975; Crow and Harmeling-van der Wel, 2008): A standardized, performance-based measure of upper-limb motor function with high inter-rater and test-retest reliability in stroke populations (Duncan et al., 1983; Hsueh et al., 2008; Platz et al., 2005); with a maximum score of 66 for typical motor function. Scores in chronic stroke are typically categorized as mild (> 45), moderate (30-45), and severe (< 30) impairment (Bushnell et al., 2015). (ii) *Box and Block Test (BBT) (*Mathiowetz et al., 1985): A measure of unilateral gross manual dexterity in stroke where participants transfer as many blocks as possible, one at a time, over a partition within 60 seconds, with performance scored as the number of blocks transferred. Normative values in healthy adults (20-80 years of age) are: 77 ± 11.6 blocks (right) and 75 ± 11.4 blocks (left) for males, and 78 ± 10.4 blocks (right) and 76 ± 9.5 blocks (left) for females (Mathiowetz et al., 1985). (iii) *Nine Hole Peg Test (NHPT)* (Kellor et al., 1971; Mathiowetz et al., 1985): A standardized measure of fine finger dexterity with excellent test-retest reliability in chronic stroke (ICC = 0.99) (Ekstrand et al., 2016), where participants place and remove nine pegs from a board as quickly as possible (with a maximum time of 60 seconds), with performance recorded as completion time. Normative values in healthy adults are approximately 19.0 ± 3.2 sec (right) and 20.6 ± 3.9 sec (left) for males, and 17.9 ± 2.8 sec (right) and 19.6 ± 3.4 sec (left) for females (Mathiowetz et al., 1985).

BBT was used as the primary measure of motor function for most analyses, as it served as the primary outcome measure in the overarching study. FMA-UE and NHPT were used along with BBT in the dimensionality reduction analysis to capture multiple aspects of motor function within a high-dimensional framework.

### Somatosensory Evoked Potential (SEP) Analysis

The EEG data was pre-processed by removing bad channels, notch filtering (55-65 Hz), and band pass filtering at 0.2 - 40 Hz (zero-phase causal digital filter– Butterworth Filter, model order 2). Ocular artifacts were removed with referential Independent Component Analysis (James and Hesse, 2005, Jiang et al., 2019), using Fp1 or Fp2 as a reference. Xdawn based spatial filtering (Rivet et al., 2009) was used to improve the signal to noise ratio of the evoked response with respect to background noise. The preprocessed data was epoched in 400 msec long segments (-100 msec to 300 msec), which were averaged to obtain the SEP response, and grand averaged across participants.

To assess across-trial spatio-temporal consistency, or the signal-to-noise ratio (SNR), of the SEP response to each stimulated hand, and its spatial topography, we computed and visualized the signed coefficient of determination (r^2^) across all channels. The r^2^ topography was visualized to assess response specificity. For visualization purposes, grand-average topographies were computed by combining responses to the affected and unaffected hands, with lateral inversion (left-right flipping) applied as needed to align hemispheric representations.

We additionally assessed a binarized representation of the SEP responses to evaluate group differences between participants with very weak versus moderate responses, complementing the continuous analysis. The threshold for binarization was defined as an SEP amplitude indistinguishable from baseline noise, representing responses that were absent (smaller or equal to baseline). This criterion provided a conservative means of distinguishing participants with absent or severely diminished cortical responses from those with measurable responses.

#### SEP Decoding

With the aim to discriminate the SEP responses from the affected versus unaffected hand stimulation, we apply a regularized linear discriminant analysis (rLDA) (Fisher 1936). LDA is a supervised machine learning algorithm that maximizes class separability by optimizing the ratio of between-class variance to within-class variance. We used the responses at centro-parietal electrodes (C3, P3, Cz, P4, C4) within 50-80 msec (root mean square) from stimulus onset as input features. Data set was balanced, with approximately 300 trials per hand. Classifier performance was evaluated with a 5-fold cross-validation method (Cecotti and Ries, 2017), where we trained the model on 80% of the data and tested on the remaining 20% in each fold. The classification was quantified using the area under the receiver operating characteristic curve (AUC). AUC values range from 0 to 1 (perfect classification), where 0.5 indicates chance level. The overall performance was calculated as the median AUC across all folds. A high AUC indicates separability of sensory representations across the two hemispheres.

Further, to assess whether the classification performance was beyond chance, we applied a permutation test. The class labels were randomly shuffled 1000 times, and for each permutation, an LDA classifier was trained on the data with the shuffled labels. The classification performance was quantified by the AUC, providing a null distribution of the AUC values at chance-level performance. The statistical significance (p-value) of the observed AUC was then determined as the proportion of the permuted AUC values greater than or equal to the true AUC.

#### Vibration-based Event Related Desynchronization (ERD) Analysis

To assess the mu band (8-12 Hz) suppression associated with vibration onset, we analyzed the preprocessed EEG data using a short-time Fourier transform (STFT) with a 130 msec Hanning window to obtain time-frequency representations. Mu-band power was extracted in the post-stimulus window of 150-300 msec relative to vibration onset, over parietal electrodes (P3/P4) contralateral to the stimulated hand. Event-related desynchronization (ERD) was quantified as the percentage change in power relative to a pre-stimulus baseline window of - 100 to 0 msec. To assess the spatial specificity of ERD, we visualized scalp topographies for stimulation of the affected and unaffected hands, averaged across participants. We then evaluated the relationship between ERD and the corresponding SEP magnitude by assessing correlations for both the affected and unaffected hands.

### t-distributed stochastic neighbor embedding (t-SNE) (van der Maaten and Hinton, 2008)

We apply the nonlinear dimensionality reduction technique t-SNE to a combination of heterogenous motor function measurements to visualize the underlying structure of high-dimensional data in a two-dimensional embedding. Four measures were included: Box and Block Test (BBT), Nine Hole Peg Test (NHPT), Fugl-Meyer Assessment of the Upper Extremity (FMA-UE), and NIH Stroke Scale (NIHSS). Prior to the analysis, the scores were curated following procedures similar to Gupta et al. (2018): (i) values were aligned such that lower scores indicated poorer function, (ii) normalized to their respective maximum possible values, and (iii) standardized using z-scores. T-SNE was applied in MATLAB to the normalized scores using standardized inputs and a perplexity of 8. The resulting two-dimensional embedding was clustered using hierarchical agglomerative clustering with average linkage and Chebyshev distance. The optimal number of clusters in the t-SNE embedding was determined using the Calinski-Harabasz index (Caliński and Harabasz, 1974), which maximizes between-cluster dispersion while minimizing within-cluster dispersion, providing a measure of clustering quality. The optimal number of clusters was selected as the value that maximized the index over a predefined range of cluster numbers (1-6), avoiding over-partitioning at higher cluster counts.

As the t-SNE is stochastic in nature, the embedding was repeated multiple times, and the consistency of cluster assignments across runs was quantified using the Adjusted Rand Index (ARI) (Hubert and Arabie, 1985, Warrens and van der Hoef 2022). The ARI quantifies similarity between two clustering outcomes by evaluating pairwise agreement, counting the number of element pairs that are consistently assigned to either the same cluster or different clusters. It ranges from -1 to 1, where 0 is considered chance level agreement and 1 indicates perfect agreement; values between 0.3 and 0.7 are considered to show moderate reproducibility.

We then evaluated whether the somatosensory measures of SEP discriminability and proprioceptive error differed significantly across the identified clusters. Finally, cluster membership was mapped onto pairwise motor and stroke severity scores to assess whether the t-SNE derived clusters corresponded to distinct functional levels.

### Statistical Analysis

Paired testing was performed with the nonparametric Wilcoxon signed-rank test, and the independent two sample test was performed with the Mann-Whitney U Test.

*P* values < 0.05 were considered significant. Correlation analysis was performed with the Spearman’s *ρ*. The effect size was calculated with a rank-based *r* (Field 2018): *r* = *Z*/√*n*, where *Z* is the test statistic, and *n* is the total number of observations (i.e., *n*_1_ + *n*_2_for the independent Mann-Whitney U Test, and number of pairs for the paired Wilcoxon signed-rank test). Repeated measurements were tested with the Friedman test (Friedman, 1937). Its effect size was estimated using the coefficient of concordance (Kendall’s (Tomczak and Tomczak, 2014)) with the equation *w* = *χ*^2^⁄*n*(*k* − 1), where *χ*^2^ is the Friedman test statistic, *n* is the sample size, and *k* is the number of repeated measurements. The interpretation of *r* and Kendall’s *w* was based on Cohen’s interpretation guidelines (Cohen 1977) of 0.1 – < 0.3 (small effect), 0.3 – < 0.5 (moderate effect), and >= 0.5 (large effect). Kruskal-Wallis test was used to determine significant differences between 3 or more independent groups. *Post hoc* analysis was performed with Mann-Whitney U tests. The effect size was calculated by *η*^2^ = (*H* − *k* + 1)/ (*n* − *k*), where *H* is the test statistic, *k* is the number of groups, and *n* is the total number of samples; with an interpretation of ∼0.01 as small, ∼0.06 as medium, and ≥ 0.14 as large. Multiple comparisons within each set of related analyses were controlled using the Benjamini–Hochberg false discovery rate (FDR) procedure.

Agreement between the two baseline proprioception measurements was assessed using the intraclass correlation coefficient (ICC) (Koo and Li, 2016; Liljequist et al., 2019). ICC estimates were computed in MATLAB R2020b using a two-way mixed-effects model for single measurements with absolute agreement (ICC (3,1), or also referred to as A-1). ICC outcomes were interpreted according to established guidelines (Koo and Li, 2016) as follows: < 0.5 (poor), 0.5-0.75 (moderate), 0.75-0.90 (good), and > 0.90 (excellent) agreement. The coefficient of determination (*r*^2^) was used to calculate the signal-to-noise ratio and the spatial specificity of the scalp regions most active during the evoked response. As this assessment is binary, the labels used are arbitrary (-1, +1), and hence a point biserial correlation coefficient was used. Classification performance was evaluated using the area under the receiver operating characteristic curve (ROC AUC). AUC values range from 0 to 1 (perfect classification), with 0.5 being chance level (Bradley, 1997).

## Results

We analyzed neurophysiological responses to finger vibrotactile stimulation and finger proprioception error in 46 people with chronic stroke. They included 12 women and 34 men who were 57 ± 15 years of age (mean ± standard deviation) and had a stroke 4.7 ± 3.5 years ago. The group included 42 participants with a minor stroke (i.e., an NIHSS stroke severity score of 0-4), and 4 participants with a moderate stroke (NIHSS score of 5, 6 or 7). Twenty participants had a hemorrhagic stroke, while 25 had an ischemic stroke; and one participant had both. Twenty-six people had a left upper limb impairment while 20 people had a right limb impairment. The first participant’s EEG data was excluded in this analysis as it was collected with an alternate EEG software setup.

The average BBT score (mean ± standard deviation: 24 ± 15.3) for the affected hand was substantially lower than normative values reported for healthy adults (∼75-78 blocks), indicating marked UE impairment. The average NHPT score for affected hand was 53.7 ± 9.8 sec (task time was capped at 60 sec), which was also markedly higher than healthy adults (∼18-20 sec). The average FMA-UE score for the affected hand was 45.9 ± 11.1 out a possible total of 66, indicating moderate impairment severity across the group. Proprioception data were collected at two separate baseline sessions, that were found to be in moderate agreement as assessed by ICC (ICC (3,1) = 0.74, 95% CI [0.56, 0.85], *p* = 6.1e-09), and therefore averaged to reduce measurement variability, and to obtain a single, more reliable estimate of proprioceptive performance. The average proprioception error was 12.9 ± 6.3 degrees, also markedly larger than that observed in healthy adults (2.6 ± 6.3 degree) (Ingemanson et al., 2016). In the analysis of the EEG-based somatosensory evoked responses, a mean of 298 ± 6 trials per hand per participant were analyzed after bad trials were removed. For further analysis we focus on contralateral centro-parietal scalp regions overlying the sensorimotor cortex associated with the upper limbs (Roux et al., 2018; Penfield and Rasmussen, 1950).

The aims of this study were to (i) characterize sensory neural responses to passive vibrotactile stimulation of the affected and less-affected hands after stroke, (ii) assess their relationship to oscillatory dynamics, (iii) quantify the ability to discriminate the location of sensory stimulation via decoding of bi-hemispheric responses to affected versus less-affected hand stimulation, and (iv) determine their associations with proprioceptive and motor function.

Hereafter, the term ‘unaffected’ is used to refer to the less-affected side, mainly for readability, while acknowledging that the ipsilesional limb may not be entirely unimpaired.

### Reduced Contralateral SEP Responses for the Affected Hand

Across participants, SEP-prominent peak observed around 50-75 msec (Figure 2a) recorded over the contralateral parietal cortex, was larger on average, for the unaffected hand compared to the affected hand. This likely corresponds to the well-known P50 component. Representative topographies within that segment (70 ± 5 msec) are shown in Figure 2c. They show a spatially specific contralateral parietal activation in response to stimulation of the unaffected hand, and a weaker, though still spatially localized, response for the stimulation of the affected hand. The SNR of the response, as measured by r^2^, shown as a time-domain signal in Figure 2b, was also markedly lower for the affected hand, particularly around the latency corresponding to the SEP P50 component. Based on Friedman repeated measures test, this contralateral SEP P50 of the affected hand was found to be significantly smaller than that for the affected hand (*χ*^2^= 9.8, *p* = 0.002, *w* = 0.22) (Figure 3a).

**Figure 2:**
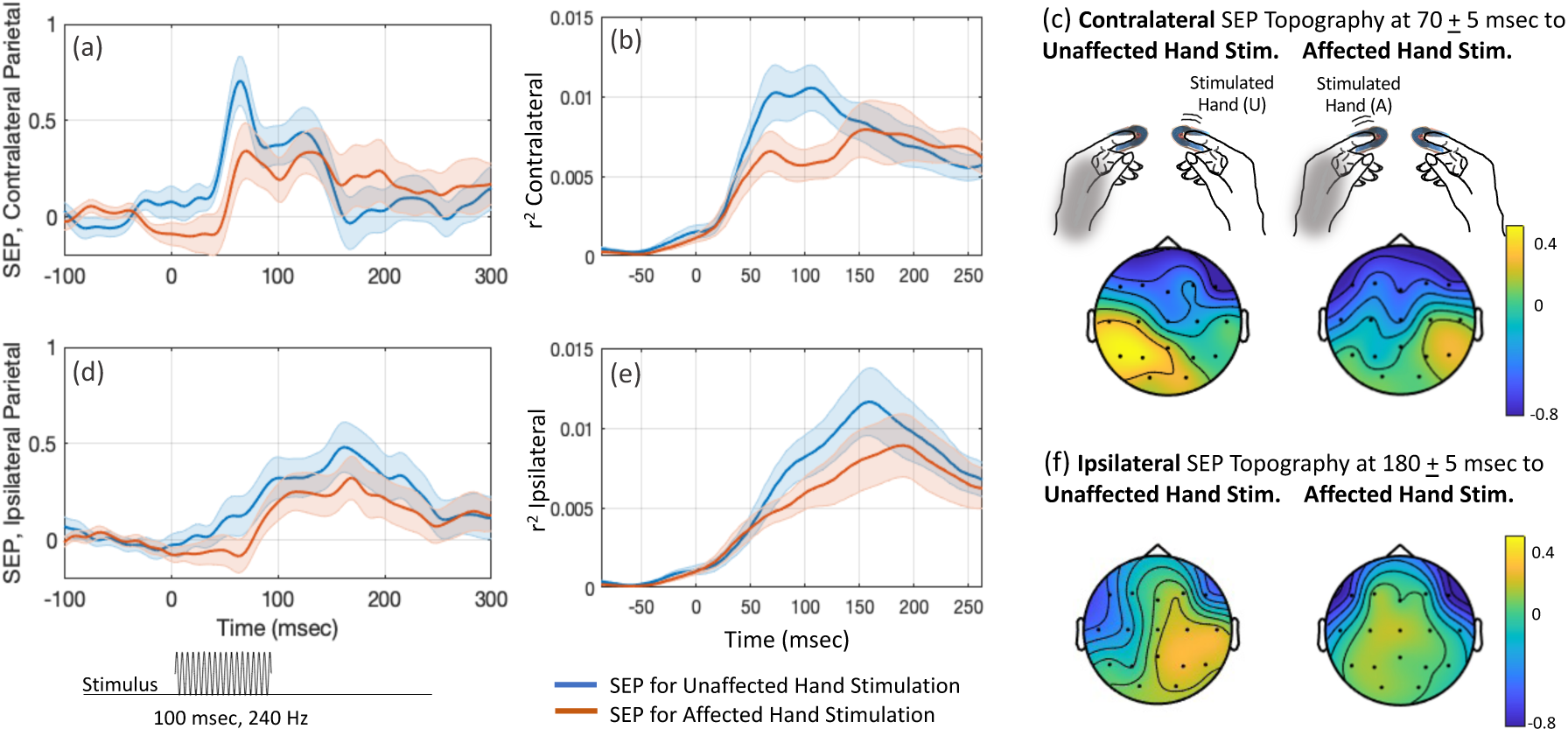
Somatosensory Evoked Potentials (SEPs) recorded over the parietal cortex, evoked by vibrotactile stimulation of the unaffected and affected hands in people with stroke: (a) and (d) show the contralateral and ipsilateral time-domain SEP signals evoked by stimulation of the unaffected hand (blue) and the affected hand (rust), respectively. The stimulation duration is 100 msec relative to onset at 0 msec. (b) and (e) show the corresponding time-domain signals for the coefficient of determination (r^2^). (c) and (f) show the scalp topographies of the evoked responses at prominent peaks: 65-75 msec for the contralateral response and 175-185 msec for the ipsilateral response, for stimulation of the unaffected and affected hands, respectively.

**Figure 3:**
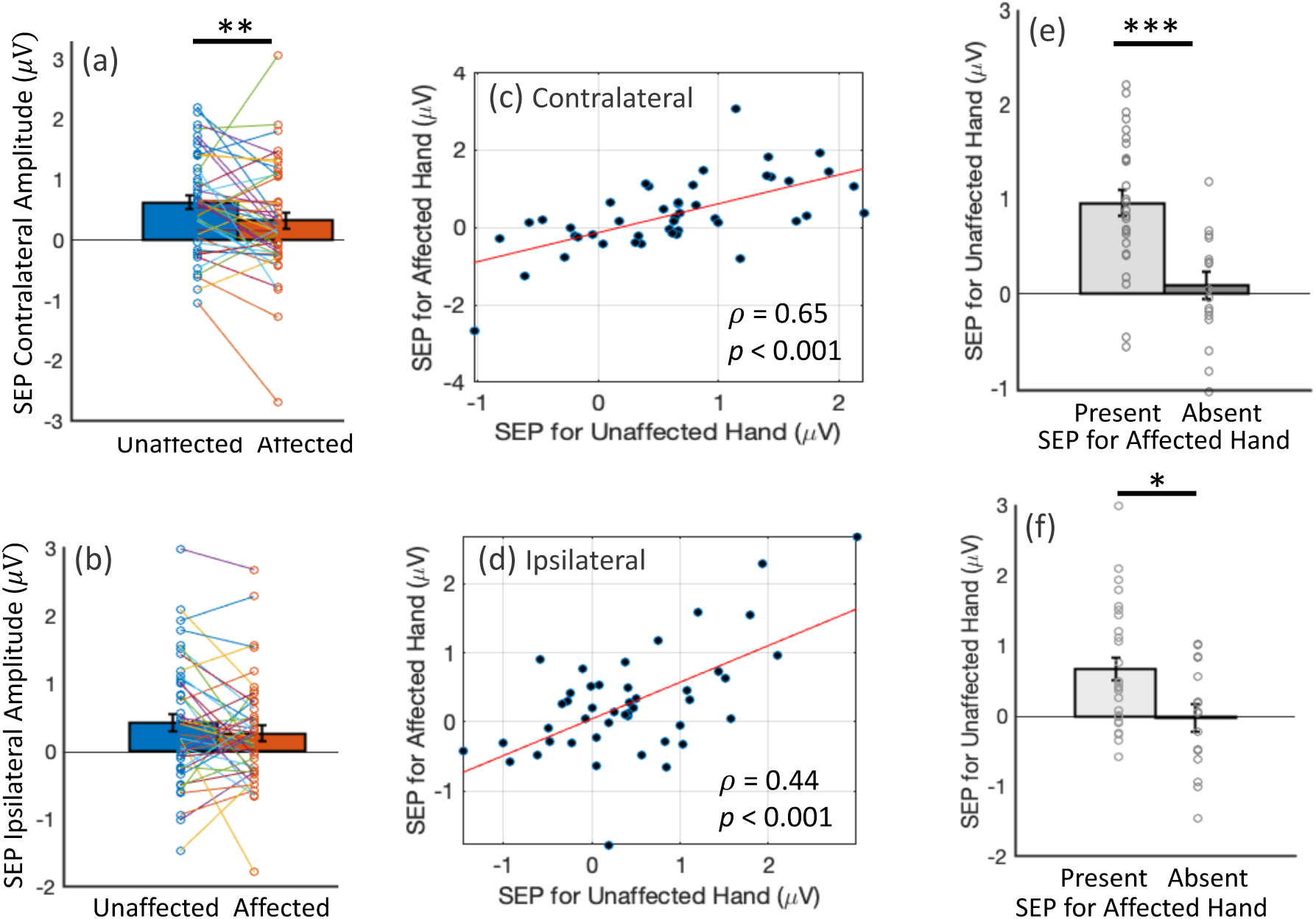
Association of somatosensory evoked potentials (SEPs) recorded over the parietal cortex, evoked by vibrotactile stimulation of the unaffected and affected hands in people with stroke: (a) and (b) show the paired associations for contralateral and ipsilateral SEP magnitude between the two hands, with a significant difference at the contralateral cortex (p < 0.01, Wilcoxon Signed-rank test). (c) and (d) show significant correlations between SEPs evoked by stimulation of the unaffected and affected hands at the contralateral cortex (at 65-75 msec, ρ = 0.65, p < 0.001) as well as the ipsilateral cortex (at 175-185 msec, ρ = 0.44, p < 0.001). (e) and (f) show a binarized association: individuals with an absent SEP for the affected hand show significantly smaller SEPs for the unaffected hand. This effect was significant in both the contralateral (p < 0.001) and ipsilateral (p < 0.05) cortex, using the Mann-Whitney U test.

At the ipsilateral parietal cortex, we observed a late positivity at approximately 175-185 msec, with similar morphology for both unaffected and affected hand stimulation (Figure 2d). Representative topography in that segment (180 ± 5 msec) (see Figure 2f) showed a greater spatial specificity for the unaffected hand. The corresponding r^2^ peaked at that latency, for both hands, indicating comparable SNR (Figure 2e). Supporting the above observations, the ipsilateral SEP response for the affected hand was not found to be statistically significantly different (*χ*^2^= 0.56, *p* = 0.456) from that of the unaffected hand (Figure 3b).

### Preserved Interlimb Association of SEP Responses

Next, we assessed whether SEP responses for the affected hand were associated with those of the unaffected hand across participants. The contralateral SEP P50 elicited by the stimulation of the affected hand was strongly correlated with the corresponding response from the unaffected hand (*ρ* = 0.6, *p* = 1.05 e-06) (Figure 3c). To further evaluate this relationship, a binary representation of the affected-hand SEP response was analyzed, categorizing the response as absent (amplitude less than or equal to baseline) or present. This binarization highlights a subgroup of participants with very weak responses. Participants with *absent* SEP P50 responses in the affected hand showed significantly smaller SEP amplitudes for the unaffected hand compared to those with present responses (*p* = 7.817e-05, *r* = 0.56, Mann-Whitney U test, Figure 3e).

A similar analysis of the late ipsilateral SEP response (175-185 msec) showed a weaker but significant correlation between the affected and unaffected hands (*ρ* = 0.44, *p* = 0.001) (Figure 3d). And even though no significant difference was observed between the ipsilateral SEP magnitudes of affected and unaffected hands in the continuous analysis (Figure 3b), the binarized representation showed a modest but significant group difference (*p* < 0.05, *r* = 0.34, Mann-Whitney U test) (Figure 3f).

### SEP Magnitude Relates to Oscillatory Sensorimotor Activity

Spectral analysis of the oscillatory sensorimotor activity over the centro-parietal cortex showed event related desynchronization (ERD) occurring at 150-300 msec following the onset of vibrotactile stimulation. This activity was observed for both affected and unaffected hands, with a greater magnitude (expressed as a percentage of baseline sensorimotor rhythm) and a more spatially central distribution, as shown in Figure 4a. The ERD over the parietal cortex induced by the stimulation of the affected hand was significantly reduced compared to the unaffected hand (*p* = 0.037, Mann-Whitney U test, Figure 4b). Further, ERD magnitude was significantly correlated with the corresponding contralateral SEP P50 magnitudes, both for the affected hand (*ρ* = 0.36, *p* = 0.007) and the unaffected hand (*ρ* = 0.35, *p* = 0.008). This association indicates that early sensory responses and subsequent oscillatory activity may reflect coupled mechanisms of cortical sensorimotor processing.

**Figure 4:**
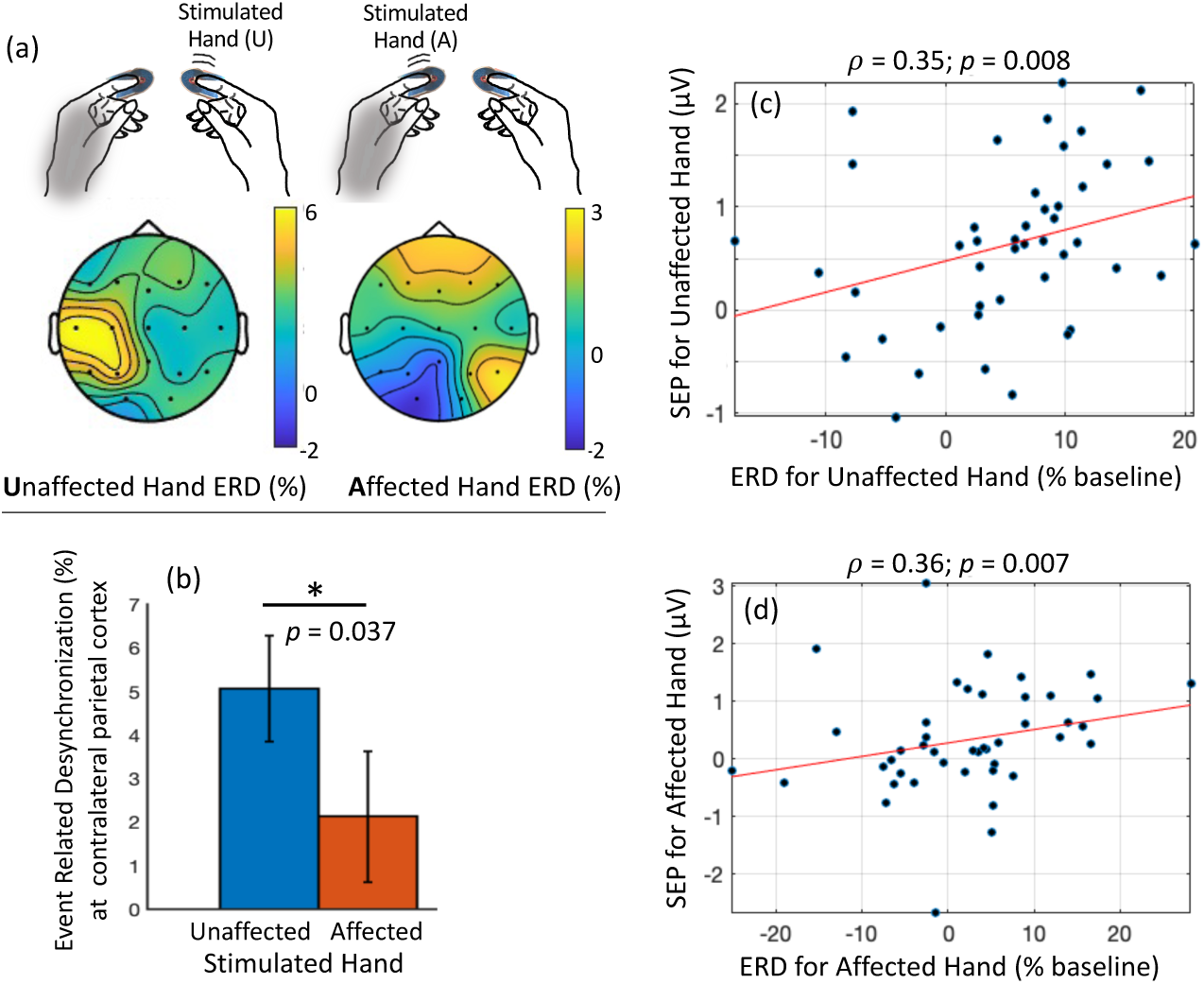
Event Related Desynchronization (ERD) induced by vibrotactile stimulation and its association with somatosensory evoked potentials (SEP) in people with stroke: (a) Scalp topographies illustrating the spatial distribution of ERD (percentage of baseline activity) occurring 150–300 msec following stimulation onset for the unaffected and affected hands. (b) ERD over the parietal cortex was significantly reduced for the affected hand (red) compared to the unaffected hand (blue) (Mann–Whitney U test, p = 0.037). (c–d) ERD magnitude was significantly correlated with the corresponding contralateral SEP P50 amplitude for both the unaffected hand (ρ= 0.35, p= 0.008) and the affected hand (ρ= 0.36, p= 0.007).

### Relationship of Ipsilesional Sensory Responses with Motor and Proprioceptive Ability

To evaluate the association between ipsilesional (contralateral) SEP P50 magnitude and oscillatory response (ERD) and functional outcomes of the affected hand, we examined their relationship with motor and proprioceptive measures. The contralateral SEP magnitude at the injured cortex was not significantly correlated with BBT score (BBT: *ρ = -*0.01*, p =* 0.467). Similarly, contralateral vibration-induced ERD magnitude was not significantly correlated with BBT score (*ρ* = -0.02, *p* = 0.452).

In contrast, this SEP magnitude was significantly correlated with proprioceptive error (*ρ* = -0.32, *p* = 0.03, Figure 5a), with lower SEP magnitudes being associated with greater proprioceptive impairment. Using the binarized SEP classification described above, participants with absent SEP responses exhibited significantly greater proprioceptive error compared to those with present responses (Figure 5b, *p*= 0.030, *r* = 0.31, Mann-Whitney U test). However, ERD was not found to be correlated with proprioceptive error (*ρ* = -0.12, *p* = 0.227).

**Figure 5:**
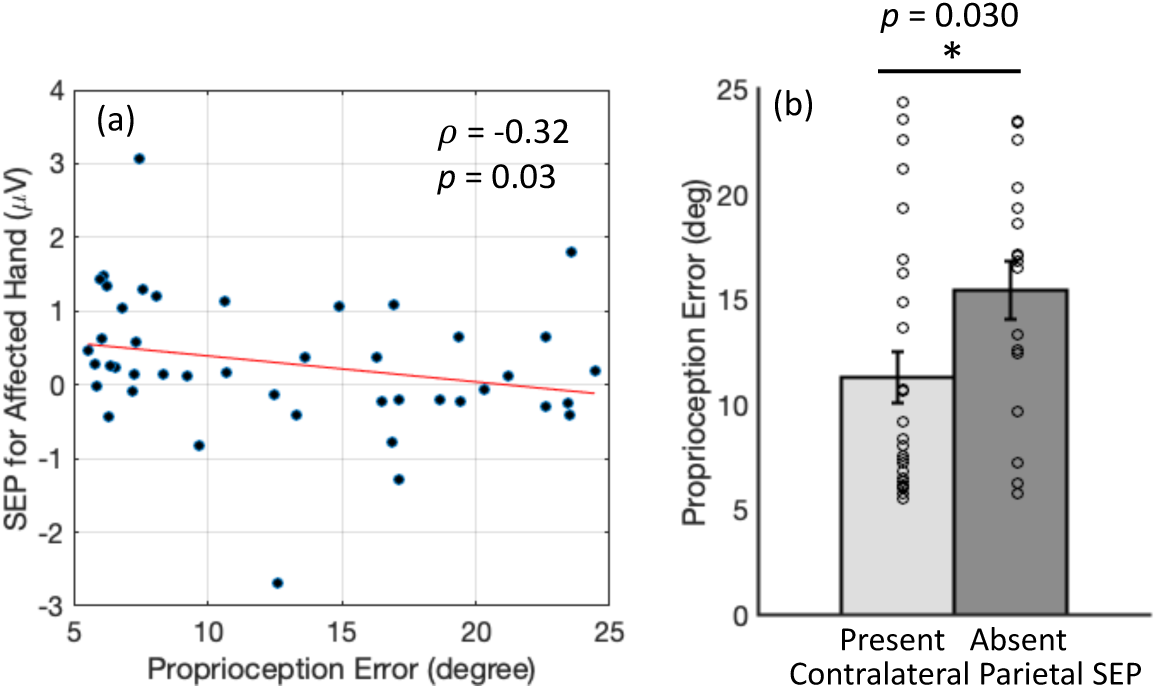
Association between vibrotactile somatosensory evoked potentials (SEPs) and proprioception errors measured with the Finger Individuating Grasp Exercise Robot (FINGER): (a) Significant correlation between the contralateral SEP P50 evoked by stimulation of the affected hand and proprioception error (in degrees) measured at the affected hand (ρ = -0.32, p = 0.03). (b) Binarized comparison of the same measures, based on the presence or absence of contralateral SEPs evoked by stimulation of the affected hand. Participants with indetectable (absent) SEPs exhibited significantly larger proprioceptive errors (p = 0.03, Mann-Whitney U test).

### Bihemispheric Decoding of Somatosensory Responses is Associated with Both Proprioception and Motor Function

In contrast to the analysis of SEP magnitude in the injured parietal cortex contralateral to the affected hand, which reflects local sensory processing, we next evaluated the *discriminability* of SEP responses. Here, discriminability refers to the extent to which a decoder can identify the stimulated hand (affected vs. unaffected) from the combined bihemispheric SEP responses, thereby capturing integrated sensory processing across both hemispheres. Notably, this analysis captures trial-by-trial variability, including the consistency of responses within each condition (stimulation of a hand), as well as the separability between conditions (stimulation of affected vs unaffected hand), which may not be apparent in trial-averaged waveforms. SEP decoding was performed with an rLDA, with five-fold cross-validation, and the classification performance was quantified using the area under the receiver operating characteristic curve (AUC).

Mean AUC scores were 0.77 ± 0.09. Permutation testing with shuffled labels confirmed that this classification performance was significantly above chance for all participants (all p < 0.001). Subject-wise AUC score was significantly correlated with proprioceptive error at the affected hand (*ρ* = -0.39, *p* = 0.004, Figure 6a), with higher SEP discriminability associated with a smaller proprioceptive error.

**Figure 6:**
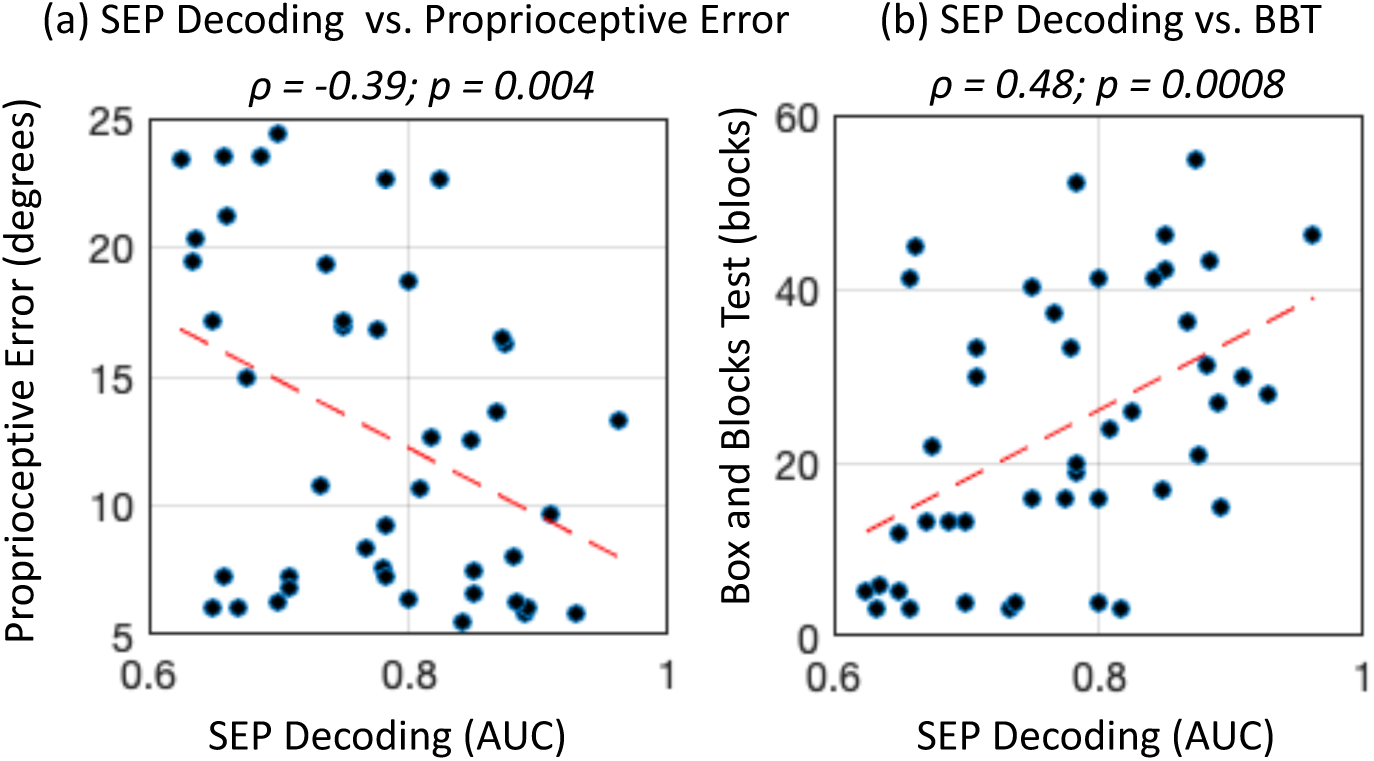
Correlation between somatosensory evoked potential (SEP) discriminability and proprioceptive and motor function: SEP decoding performance was significantly correlated with (a) proprioceptive error (ρ = -0.39, p = 0.008), (b) Box and Block test (BBT) scores (ρ = 0.48, p = 0.0008).

Importantly, the bihemispheric SEP classification performance was also significantly correlated with motor function at the affected hand, as measured by the BBT (*ρ* = 0.48*, p* = 0.0008, Figure 6b). Linear regression showed that SEP decoding explained approximately 22.5% of the variance in baseline motor function (*β* = 79.8, *r*^2^= 0.225, *p* = 0.001). These results indicate that a greater neural discriminability – i.e., better somatosensory classification – is associated with both better proprioceptive and motor performance at the affected hand.

### SEP Discriminability Differs Across Subgroups Identified by t-SNE-Based Joint Analysis of Motor Function

Finally, we performed a joint analysis of sensorimotor performance across clinical stroke severity and motor function assessments, capturing complementary aspects of motor ability, including gross and fine finger dexterity, as well as inherent somatosensory processing. The aim was to determine whether distinct subgroups emerged from this multidimensional representation and whether such subgrouping could be explained by SEP discriminability, thereby highlighting a potentially obscured sensory component.

The resulting two-dimensional t-SNE embedding revealed three distinct clusters (Figure 7a). The Calinski-Harabasz index identified three as the optimal number of clusters, with the index value of 130. Cluster stability was assessed across 190 pairwise t-SNE reruns, with cluster similarity quantified using the Adjusted Rand Index (ARI). The mean ARI was 0.6 ± 0.2, indicating moderate consistency of the embedding and clustering results.

**Figure 7:**
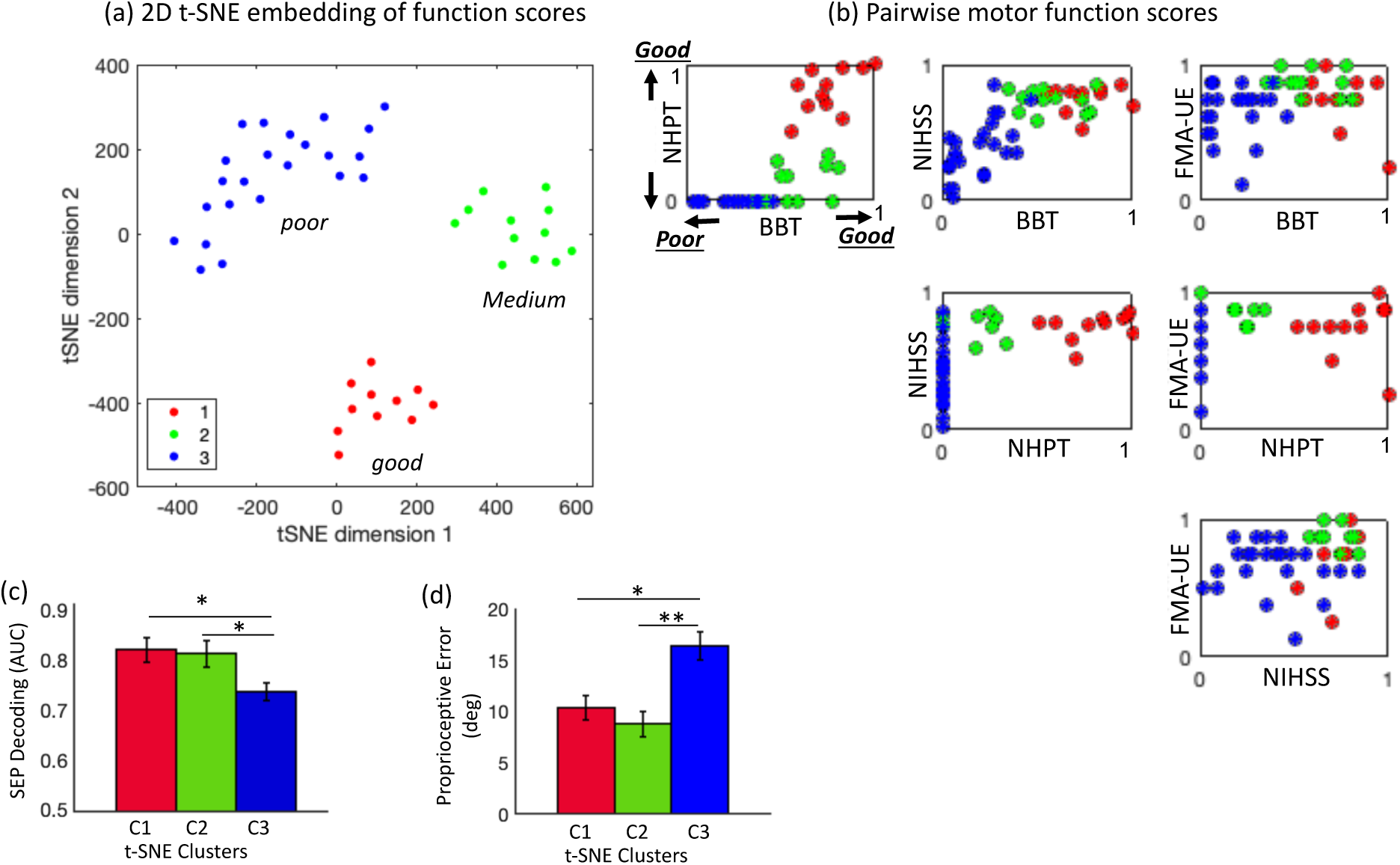
t-distributed stochastic neighbor embedding (t-SNE)-based nonlinear dimensionality reduction of motor function measures: (a) Two-dimensional t-SNE embedding shows 3 distinct and moderately stable clusters (Adjusted Rand Index: 0.6 ± 0.2). (b) Mapping the cluster membership onto pairwise motor function plots shows correspondence with poor, medium and good function levels. (c) Somatosensory evoked potential (SEP) decoding performance, quantified by the Area Under the Curve (AUC), differs significantly across clusters (p = 0.014, Kruskal-Wallis test), with a large effect size (η^2^= 0.15). (d) Proprioceptive error (degrees) also differs significantly across the clusters (p = 0.003, Kruskal-Wallis test) with a large effect size (η^2^ = 0.22).

The mean SEP discriminability (AUC) across clusters was 0.82 ± 0.08 (cluster 1, red), 0.81 ± 0.09 (cluster 2, green), and 0.73 ± 0.08 (cluster 3, blue) (Figure 7a). These differences were statistically significant (H(2) = 8.46, *p* = 0.014, Kruskal-Wallis test) with a large effect size (*η*^2^ = 0.15) (Figure 7c). *Post hoc* analysis showed a significant difference between clusters 1 and 3 (*p* = 0.017, r = 0.41) and clusters 2 and 3 (*p* = 0.021, r = 0.39), both indicating moderate effect sizes. When aligned with pairwise motor assessment scores (Figure 7b), the identified clusters corresponded to poor (cluster 3), moderate (cluster 2), and good (cluster 1) performance levels. SEP discriminability tended to increase with motor function, with the lowest, intermediate, and highest discriminability corresponding to poor, moderate, and good motor performance, respectively (Figure 7c).

These clusters also showed a similar trend for proprioceptive error. The mean proprioceptive error across clusters was 10.35 ± 4.0 (cluster 1, red), 8.78 ± 4.4 (cluster 2, green), and 16.38 ± 6.2 (cluster 3, blue) (Figure 7d). These differences were also significantly different (H (2) = 11.2, p = 0.003, Kruskal-Wallis test) with a large effect size (*η*^2^ = 0.22). *Post hoc* analysis showed a significant difference between clusters 1 and 3 (*p* = 0.015, r = 0.42), and clusters 2 and 3 (*p* = 0.005, r = 0.48). The proprioceptive error followed a similar functional stratification– with the highest error observed in cluster 3 (poor motor function) and lower errors in clusters 1 and 2 (moderate and good motor function, respectively). The contralateral SEP magnitude at parietal cortex and ERD magnitude were not found to be significantly different across clusters (H (2) = 0.32, *p* = 0. 853) and (H (2) = 1.76, *p* = 0.41) respectively.

## Discussion

In this study, we characterized cortical sensory processing in forty-six participants following stroke using vibrotactile EEG-based somatosensory responses acquired with a rapid passive paradigm and portable dry-electrode EEG system. Ipsilesional sensory responses to affected hand vibration were reduced. SEP magnitude (but not ERD) as well as bihemispheric decoder performance were significantly correlated with finger proprioceptive error. In contrast, only decoder performance was strongly associated with motor function of the affected hand. Furthermore, nonlinear subgrouping analysis identified distinct functional subgroups associated with differences in cortical sensory discriminability. Together, these findings suggest that distributed sensory representations and interhemispheric sensory discriminability provide functionally relevant markers of sensorimotor integrity following stroke, and therefore may support future sensory-driven rehabilitation (Reinsdorf, et al., 2011, Sasaki et al., 2025, Bolognini et al., 2016, Carey et al., 2011, Schabrun and Hillier, 2009) and patient stratification approaches.

### Impaired Ipsilesional Sensory Processing and its Relationship with Oscillatory Sensorimotor Dynamics

Evoked (SEP) and induced (ERD) somatosensory responses to stimulation of the affected hand were significantly reduced at the contralateral (ipsilesional) cortex compared with responses to the less-affected hand at the contralesional cortex. This is in line with previous studies where sensory responses at the injured cortex have been shown to be attenuated, reflecting direct or indirect disruption in the cortical and/or subcortical sensory connections, as opposed to a peripheral or spinal injury (Sahrizan et al., 2025). This has been demonstrated by multiple neuroimaging studies, both physiological and anatomical (MEG, EEG, fMRI, MRI) at multiple spatial and temporal resolutions. Findings include diminished cortical evoked potential amplitudes and delayed responses (Eksantivongs et al., 1991), reduced ipsilesional primary somatosensory cortex activation (Carey et al., 2006), disrupted thalamocortical connectivity (Grefkes et al., 2008), and structural degeneration of sensory pathways (Schaechter et al., 2009). Some of these alterations have also been shown to be associated with impaired sensory function (Carey, 2017, Connell et al., 2008, Gupta et al., 2017).

ERD, or mu suppression, induced by vibration, is also a useful measure to characterize the oscillatory dynamics involved in sensory processing (Chung et al., 2013, Kim et al., 2020). It is commonly interpreted as a marker of cortical engagement during sensory processing, with the mu rhythm representing an idling state of sensorimotor networks (Pfurtscheller et al., 1996, Pfurtscheller and da Silva, 1999). Although this same rhythm has been extensively studied in relation to movement (Pfurtscheller, 2001), vibration-induced mu suppression occurs following sensory input and may reflect cortical readiness for integrating incoming sensory information (Kuhlman 1978). In contrast, movement-related mu suppression, which typically precedes or accompanies movement initiation, is thought to reflect cortical readiness for motor execution (Pfurtscheller, 2001, Pfurtscheller et al., 1997). While these processes may be related, their relationship was not directly assessed in the present study and may be useful to explore in future work.

SEP captures phase-locked evoked responses while ERD measures non-phase-locked oscillatory modulation (ERD or mu suppression). These measures were modestly correlated with each other at both hemispheres, a finding that to our knowledge has not been reported previously. SEP magnitude has been shown previously to be associated with pre-stimulus oscillatory state (Jones et al., 2010, Iemi et al., 2019, Linkenkaer-Hansen et al., 2004, Ritter and Becker 2009) and has often been examined in relation to trial-to-trial variability. Here, the association between SEP magnitude and vibration-induced change in oscillatory activity (ERD) suggests that these measures reflect related underlying sensorimotor mechanisms. Specifically, individuals with larger ERD showed larger SEP responses, consistent with greater cortical engagement and more effective sensory transmission, respectively.

In this study, we used vibrotactile stimulation instead of the more commonly used electrical stimulation as it provides more naturalistic activation of the somatosensory pathways. We use a vibration frequency of 240 Hz to preferentially stimulate Pacinian corpuscles (fast-adapting type II afferents) located in the dermal layers (Johnson et al., 2000, Johnson 2001). These mechanoreceptors are primarily responsible for detecting high-frequency vibrations, with peak sensitivity around 250 Hz. Multiple studies have shown that stimulation in the frequency range of 175-250 Hz robustly activate somatosensory cortical pathways (Rustamov et al., 2025, Mountcastle et al., 1967, Johnson 2001, Gupta et al., 2017, Harrington and Downs, 2001). They are involved in detecting fine textures and contributing to object perception. As the Pacinian corpuscles are densely concentrated in the finger pads (Germann et al., 2021), they are particularly relevant for studying sensory function related to dexterous hand use. Even though vibrotactile SEPs typically have lower SNR compared to electrically evoked SEPs and may be more sensitive to fluctuations in arousal, attention, or fatigue, they pass through natural mechanoreception transduction and can be more representative of functional tactile processing. To mitigate the lower SNR, we presented a considerable number of stimuli (300 per hand) in a rapid paradigm. In addition, data-driven approaches, including independent component analysis (James and Hesse, 2005), spatial filtering (McFarland 2015), and xDAWN (Rivet et al., 2009), were used to retrospectively denoise the data and improve signal quality.

### Ipsilesional SEP Responses are Associated with Proprioceptive Function

Motor control relies on cutaneous mechanoreceptive and proprioceptive feedback. Cutaneous feedback is captured by mechanoreceptors (e.g. Pacinian corpuscles, Merkel cells) located in the dermal and subdermal layers, that encode tactile and pressure-related stimuli at the skin surface, enabling force and grip control according to object texture and shape (Johnson et al., 2000, Johnson 2001). Whereas proprioceptive feedback involves muscle spindles and Golgi tendon organs (Sherrington, 1907; Proske & Gandevia, 2012; Proske, 2015), it includes substantial contributions from cutaneous mechanoreceptors SA-II afferents that encode skin stretch (Cordo et al., 2011, Gandevia et al., 1983, Strzalkowski et al., 2017), providing information about limb position, movement, and force. Thus, although cutaneous mechanoreception and proprioception engage different afferent receptors (FA-II rapidly adapting and SA-II slow adapting, respectively), both modalities depend on DCML integrity and fast thalamocortical transmission, through large-diameter afferents to support timely sensorimotor feedback. Consequently, the SEP-proprioception association is more likely to reflect damage to these common afferent pathways or cortical connections rather than selective activation of these peripheral receptors.

Results showed that the proprioceptive error of the affected hand was significantly correlated with the corresponding ipsilesional SEP magnitude, a stimulus-locked sensory response, but not ERD, a response that captures general cortical engagement. Further, in a binarized assessment, individuals with absent SEP responses performed significantly worse on the proprioceptive task. These observations support convergence of mechanoreceptive and proprioceptive processing within shared DCML and thalamocortical pathways. This raises the possibility that recovery of these sensory modalities may occur in parallel during rehabilitation, and that improvements in one modality may potentially facilitate the other through shared sensorimotor mechanisms, although this requires further longitudinal investigation.

We also evaluated the relationship between ipsilesional (contralateral) sensory responses–both evoked (SEP) and induced (ERD)– and motor function. Neither measure was significantly associated with motor function (BBT). Although mechanoreceptive pathways predominantly project contralaterally (Brodmann area 3b) (Hlushchuk and Hari, 2006), ipsilateral cortical responses in area 3b of the contralesional hemisphere are also known to occur, and are likely altered after injury, due to changes in the interhemispheric interactions (Allison e al., 1989, Lipton et al., 2006, Hlushchuk and Hari, 2006). We explored this further with the decoding analysis. From an analytical perspective, averaged evoked and induced responses may also be sensitive to trial-to-trial variability and fluctuations in background EEG due to arousal, attention, or fatigue. Such averaging may obscure sensory processing features that are better captured through trial-level analyses, explored with SEP decoding in this study.

### Coupled Bilateral Sensory Responses Reflect Shared Network

Our results showed that the contralateral SEP responses evoked by the stimulation of the affected and less-affected hands were significantly correlated. The binary analysis further showed that individuals with a preserved SEP response for the affected hand also had a larger SEP response to stimulation of the less-affected hand compared to individuals without detectable affected-hand responses. This suggests that the contralesional hemisphere (contralateral to the less-affected hand) was not always functionally spared, which has also been discussed in previous studies (review: Kitsos, et al., 2013). Moreover, the reduced sensory responsiveness at the ipsilesional hemisphere (contralateral to the affected hand) was accompanied by a corresponding, although smaller, reduction in contralesional response. Together, these results indicate a potential coupling (direct or indirect) of somatosensory processing at the bilateral parietal regions. The ipsilateral SEP responses also demonstrated significant interhemispheric association and binary pattern. These results motivate us to consider bilateral sensory network responses, rather than focusing on the typical dominant unilateral contralateral responses.

### Sensory Discriminability as a Marker of Integrated Sensorimotor Function

Typically, the somatosensory responses evoked by each hand are known to have a dominant presence at the somatosensory hemisphere contralateral to the stimulated hand, attributed to the crossed afferent pathways. Small ipsilateral responses have also been observed (Iwamura et al., 1994), considered to be suppressed due to reciprocal interhemispheric inhibition or deactivation (Hlushchuk and Hari, 2006, Lipton et al., 2006). Similar ipsilateral deactivation is observed for the ipsilateral primary motor cortex (M1) (Allison et al., 2000), posited to be due to corticocortical connections between somatosensory cortical area 3b and motor cortex M1 (Yumiya and Ghez, 1984). The relative magnitude of responses across the two hemispheres may therefore provide an important neural cue for identifying which hand receives the stimulation and thereby facilitating appropriate direction of the motor command.

After a stroke, however, these interhemispheric interactions and inhibitory balance can become altered due to compensatory neuroplasticity (Arya et al., 2024, Buetefisch 2015), affecting both ipsilateral and contralateral sensory representations. Consequently, sensory responses elicited by the less-affected hand at the contralesional hemisphere – which may also weaken due to global stroke-related effects – may become less distinguishable from ipsilateral responses elicited by the affected hand, thereby reducing the separability of sensory representations for the two hands. Functionally, changes in this reciprocal interhemispheric inhibition have previously been shown to affect tactile sensation (Seyal et al., 1995) and may potentially influence motor function.

To assess the relationship of altered bihemispheric sensory network responses with sensorimotor function, we quantified the decoding accuracy with which the stimulated hand (left vs. right hand) could be identified from the combined SEP responses at the two sensorimotor cortices. We used a regularized linear discriminant analysis approach, a widely used method for robust single-trial EEG-based discrimination in BCI applications (Friedman 1989, Lotte et al., 2007, Blankertz et al., 2008). The resulting SEP decoding was found to be significantly associated with proprioceptive ability but also motor function (measured by BBT). Individuals with a larger decoding accuracy showed a smaller proprioceptive error, and a higher motor function score.

This analysis allowed us to move beyond predefined spatial assumptions of lateralization (e.g., contralateral dominance) and instead assessed the discriminability of SEP responses across both hemispheres. By including a larger spatial region–spatially filtered centro-parietal regions of both hemispheres along with central Cz electrode– it may also capture within-hemispheric spatial alterations or dispersion arising from anatomical and functional reorganization after injury (Cheng et al., 2019).

Here, higher decoding accuracy reflects greater residual distinctiveness between responses evoked by stimulation of the affected and less affected hand despite injury-related damage and post-injury adaptations. This distinctiveness may facilitate more accurate attribution of the sensory response to the stimulated hand and consequently, more appropriately directed motor control, providing insight into the *functional effectiveness of the distributed sensorimotor network*, beyond physiological pathway integrity. Unlike analyses based solely on contralateral response magnitude, this approach can also capture cases in which the contralateral response remains relatively preserved but the ipsilateral response is maladaptively increased because of interhemispheric imbalance or transcallosal pathway damage, thereby reducing the distinctiveness between responses evoked by the two hands.

Additionally, unlike trial-averaged measures, decoding also incorporates the assessment of trial-to-trial response consistency, which can be affected by fluctuating arousal, attention or fatigue, commonly experienced post stroke. In an averaged response measurement, these fluctuations can distort the resulting averaged response. Such neurophysiological changes can be quite heterogeneous across individuals, depending on the extent, location, and time since injury (Sahrizan et al., 2025), making them challenging to discern or predict, particularly using behavioral measures alone. In this context, a trial-by-trial decoding of sensory responses provided a more sensitive and objective index of underlying sensorimotor network integrity.

The well-understood dependence of motor control on somatosensory feedback motivates the possibility that improving sensory processing and sensory discriminability through targeted sensory rehabilitation (Sasaki et al., 2025, Bolognini et al., 2016, Carey et al., 2011, Schabrun and Hillier, 2009, Winter et al., 2022, Seo et al., 2023) may also facilitate motor recovery. However, the observed relationship does not necessarily imply that all individuals with poor motor function will have an underlying cortical sensory impairment or would equally benefit from sensory-focused rehabilitation. Despite a relatively stronger correlation, compared to contralateral response alone, we observe variability in the data, with some participants exhibiting poor motor performance despite relatively preserved SEP decoding. This may be explained by the multiple factors underlying poor motor function. In addition, various motor function assessments also integrate multiple sensory and motor processes and compensatory strategies in ways that are difficult to dissociate from a single motor assessment. A continuous motor function score may not provide a clear threshold at which cortical sensory impairment becomes functionally relevant. Joint analysis of several motor assessment outcomes may help to identify underlying functional subgroups, which motivates the subsequent joint data analysis.

### Neural Sensory Discriminability Reveals Functional Subgroups After Stroke

The joint analysis of multiple motor scores involved analyzing them as a high-dimensional dataset with t-SNE-based dimensionality reduction (van der Maaten and Hinton, 2008). The aim was to determine whether we could isolate distinct underlying subgroups and to evaluate whether these subgroupings showed the sensory processing (based on sensory decoding and proprioception) as a potential contributor.

Our results show that t-SNE analysis identified distinct functional subgroups. This was supported by the Adjusted Rand Index (Hubert and Arabie, 1985, Warrens and van der Hoef 2022), indicating that the two-dimensional embedding and resulting cluster assignments were stable and reproducible.

Furthermore, the Calinski-Harabasz index (Caliński and Harabasz, 1974) showed that the data was optimally partitioned into 3 clusters. Although each motor assessment shows a continuous distribution of impairment, their joint representation revealed distinct sensorimotor subgroups. The association between subgroup membership and sensory decoding and proprioception suggests that distributed cortical sensory processing and proprioception likely contribute to the organization of these subgroups.

Based on pairwise motor assessments, these subgroups visually aligned with a continuum of poor, intermediate and good motor function. The clusters provided a potential demarcation of the functional abilities. The SEP decoding corresponded to the level of functional impairment. The subgroups also displayed significant differences in proprioceptive function, aligned with their functional impairment, further supporting the extent of sensory deficits across the identified subgroups.

These findings demonstrate a potential approach for stratifying individuals following stroke into functionally distinct subgroups and identifying those with cortical sensory impairments using combined motor function assessments. Such stratification may facilitate more personalized rehabilitation approaches and improve overall therapeutic outcomes, although this possibility requires systematic validation in future studies.

## Limitations

A limitation of this study is that the cohort consisted of individuals with mild to moderate chronic stroke who retained residual hand function (Box and Block Test > 3). Most participants were also able to perceive the vibrotactile stimulation, as confirmed verbally. We did not evaluate individuals with more severe sensory and motor impairments, in whom behavioral perception may be further compromised by stroke-related deficits while cortical somatosensory responses may remain measurable. In such cases, SEP-based measures may provide a useful complementary approach for assessing sensory pathway integrity when behavioral assessments are difficult to administer or lack sensitivity. Future studies should evaluate the utility of these neurophysiological measures across a broader range of stroke severity.

## Conclusion

In conclusion, a rapid vibrotactile EEG paradigm acquired with a portable dry-electrode system revealed multiple cortical markers of sensory function following stroke. While reduced ipsilesional SEP magnitude was associated primarily with proprioceptive impairment, bilateral SEP discriminability captured broader aspects of sensorimotor status, demonstrating associations with both proprioceptive and motor function and identifying functionally distinct patient subgroups. These findings suggest that distributed cortical sensory representations may provide clinically meaningful information beyond traditional measures of sensory response magnitude alone. As such, EEG-based sensory biomarkers may offer a practical approach for quantifying sensory system integrity, stratifying patients, and guiding the development and application of sensory-focused rehabilitation interventions.

## Ethical Considerations

The study was approved by the local Institutional Review Board at the University of Irvine, CA. All data were collected with written informed consent. Participation was voluntary with ability to opt out at any time. The research was conducted in accordance with the principles embodied in the Declaration of Helsinki and in accordance with local IRB requirements.

## Acknowledgements

We thank all the participants for their time and participation and the UCI team for the academic collaboration.

